# fingeRNAt - a novel tool for high-throughput analysis of nucleic acid-ligand interactions

**DOI:** 10.1101/2021.12.23.474073

**Authors:** Natalia A. Szulc, Zuzanna Mackiewicz, Janusz M. Bujnicki, Filip Stefaniak

## Abstract

Computational methods play a pivotal role in drug discovery and are widely applied in virtual screening, structure optimization, and compound activity profiling. Over the last decades, almost all the attention in medicinal chemistry has been directed to protein-ligand binding, and computational tools have been created with this target in mind. With novel discoveries of functional RNAs and their possible applications, RNAs have gained considerable attention as potential drug targets. However, the availability of bioinformatics tools for nucleic acids is limited. Here, we introduce fingeRNAt - a software tool for detecting non-covalent interactions formed in complexes of nucleic acids with ligands. The program detects nine types of interactions: (i) hydrogen and (ii) halogen bonds, (iii) cation-anion, (iv) pi-cation, (v) pi-anion, (vi) pi-stacking, (vii) inorganic ion-mediated, (viii) water-mediated, and (ix) lipophilic interactions. However, the scope of detected interactions can be easily expanded using a simple plugin system. In addition, detected interactions can be visualized using the associated PyMOL plugin, which facilitates the analysis of medium-throughput molecular complexes. Interactions are also encoded and stored as a bioinformatics-friendly Structural Interaction Fingerprint (SIFt) - a binary string where the respective bit in the fingerprint is set to 1 if a particular interaction is present and to 0 otherwise. This output format, in turn, enables high-throughput analysis of interaction data using data analysis techniques. We present applications of fingeRNAt-generated interaction fingerprints for visual and computational analysis of RNA-ligand complexes, including analysis of interactions formed in experimentally determined RNA-small molecule ligand complexes deposited in the Protein Data Bank. We propose interaction-based similarity based on fingerprints as an alternative measure to RMSD to recapitulate complexes with similar interactions but different folding. We present an application of molecular fingerprints for the clustering of molecular complexes. This approach can be used to group ligands that form similar binding networks and thus have similar biological properties.

**AUTHOR SUMMARY:** We present a novel bioinformatic tool, fingeRNAt, aiming to support scientists in the analysis of complexes of nucleic acids with various types of ligands. The software automatically detects non-covalent interactions and presents them in a form that is understandable to both humans and computers. Such data can help decipher the nature of interactions between nucleic acids and ligands and determine the main factors responsible for forming such complexes in nature. fingeRNAt finds application in multiple studies, both structure- and drug discovery-oriented. Here, we analyzed the experimentally solved structures of RNA complexes with small molecules to determine which binding features are most prevalent, i.e., most common interactions or their hot spots. The results of this analysis may help elucidate the mechanisms of binding and design new active molecules. Moreover, we propose to use the data generated by our software as a new metric for the quantitative comparison of two molecule complexes. We have shown that it is more reliable than the currently used methods in certain “difficult” cases. We have shown that the results of our program can be used for high-throughput analysis of molecular complexes and the search for active molecules. We are confident that fingeRNAt will be a valuable tool for exploring the complex world of interactions of nucleic acids with ligands.

## INTRODUCTION

Nucleic acids are essential bioorganic molecules present in every living organism. Although deoxyribonucleic acid (DNA) is traditionally viewed as a mere genetic information carrier and ribonucleic acid (RNA) as a scaffold in the protein synthesis process, their functions go far beyond that (1–3). Both DNAs and RNAs regulate diverse biological pathways and thus have a central role in cellular metabolism. Non-coding DNAs constitute the majority of the human genome and regulate protein-coding sequences by acting as a binding site for other transcriptional regulatory factors, an origin of replication site, a centromere, or a telomere (4,5). Moreover, some non-coding DNAs can be transcribed into non-coding RNAs, which play a fundamental role in the cell, as they build large macromolecular machines, deliver amino acids to ribosomes, or regulate different molecular processes, e.g., by silencing genes or driving catalytic reactions.

Nucleic acids possess the ability to adopt tertiary structures and have grooves acting as binding sites for other factors. They are capable of forming complexes with other nucleic acids, proteins (6), ions (7–10), and naturally occurring small molecules, such as metabolites (11). These interactions are essential for cell functioning as they may modulate transcription and translation processes, DNA repair, splicing, apoptosis, or stress responses. Nucleic acids are also targets for synthetic small molecule drugs. DNAs remain a primary target for several anticancer chemotherapeutics (12) and potential antimicrobial compounds (13). RNA molecules, such as the bacterial ribosomes, are also known targets for a number of drugs, e.g., bacterial ribosome-targeting antibiotics (14,15). Other RNAs, such as mRNAs (16) and regulatory RNAs in humans (17), riboswitches in bacteria (18), and conserved non-coding RNAs in viruses (19), are considered as promising targets for new therapeutics (for review, see (4,20,21)).

Weak non-covalent bonds are crucial in the molecular recognition process. Their identification and characterization help elucidate the basics of intermolecular binding and support the rational design of bioactive compounds (22,23). Typically, this process requires a laborious visual inspection of three-dimensional (3D) models of complexes by structural biologists or medicinal chemists (24). With the advent of computational methods, this procedure may be supported with programs aiming at detecting and characterizing non-covalent interactions. As most of the available programs were designed to analyze protein complexes with small molecule ligands (25–29), the number of tools focusing on nucleic acids is currently very limited. LigPlot and LigPlot+ may be used to visualize nucleic acid-ligand complexes, but they display only hydrogen bonds and hydrophobic interactions (30,31). Arpeggio is a web server dedicated to detecting and visualizing interatomic interactions in protein structures; however it may be applied to DNA macromolecules (32). The recent update of the PLIP program and its web server introduced support for DNA and RNA receptors, enabling detection and visualization of several types of non-covalent bonds (33). ProLIF is a Python library developed to generate interaction fingerprints for protein, DNA, or RNA complexes from molecular dynamics simulations, experimentally determined structures, and molecular docking (34).

To facilitate high-throughput analysis of intermolecular interactions, detected non-covalent bonds might be encoded in the form of Structural Interaction Fingerprint (SIFt), which describes the existence of specific molecular interactions between all structure’s residues and a ligand. SIFt, firstly published by Deng *et al*. as a method to study protein-ligand binding, translates information about 3D interactions within the complex into a 1D binary string (bit vector). SIFt calculation consists of two main steps. First, the presence of interaction of the specified type for each residue-ligand pair is checked, and an appropriate binary value is assigned (1 if the interaction occurs and 0 otherwise). Subsequently, all calculated binary substrings are merged into one long string - SIFt, preserving the structure’s residue order. Typical SIFt applications include post-docking analyses, such as clustering molecule’s poses from molecular docking and comparing them with reference structures or scoring functions (35). It is also frequently applied in interpreting activity landscapes, supporting structural databases, and analyzing protein-ligand complexes to search for similarities, e.g., by calculating the Tanimoto coefficient of bit vectors. In rational drug discovery, SIFt supports processing virtual screening results (36,29,37) or developing new scoring functions (38,39). With the growing importance of artificial intelligence methods in drug discovery, new applications of interaction fingerprints emerged. SIFt can be associated with the information about ligand’s biological activity, thus becoming an excellent input to the machine learning algorithms. This approach was already used, e.g., in training models to predict ligands’ activity towards protein targets (40–44).

Here we present the fingeRNAt - a Python 3 program that detects and visualizes nucleic acid-ligand interactions. As an input, it takes a 3D structure of a nucleic acid (RNA or DNA) and a file containing ligands that form complexes with this macromolecule (e.g., the output from molecular docking with an external program). fingeRNAt accepts nucleic acids, small molecules, proteins, and metal cations as ligands. The output is a fingerprint - a bit vector containing information on interactions detected between interacting partners and optionally a human-readable file containing detailed information on detected interactions. By default, it detects nine interactions (see: Implementation section in Materials and Methods) but can be easily extended to detect virtually any type of interaction using a simple plugin system; the provided sample plugin file enables detection of eight additional interactions. Moreover, accompanying programs allow for convenient post-processing and visualization of detected interactions, calculation of Receptor Preferences (aka Receptor’s Interactions Hot Spots), which represent the spatial occurrence frequency of a given interaction type in receptor atoms, and Ligand Preferences (aka Ligand’s Interactions Hot Spots), which represent the spatial occurrence frequency of a given interaction type in ligand binding site.

To the best of our knowledge, there are no nucleic-acid dedicated tools for detection and classification of interactions that encode them in both machine- and human-readable formats, are highly customizable, and allow for exhaustive postprocessing such as calculation of similarity/distance metrics and interactive visualization. A detailed comparison of the fingeRNAt features (this work) and similar software tools (Arpeggio (32), PLIP2021 (33), and ProLIF (34)) can be found in S1 Table.

fingeRNAt is freely available to download from github.com/n-szulc/fingeRNAt. Program installation guide, together with an extensive manual, multiple usage examples, and Sphinx documentation, are also accessible from the repository. fingeRNAt can be used as a standalone command-line tool, but it also has an intuitive graphical user interface with the same functionalities.

## RESULTS AND DISCUSSION

fingeRNAt is a program for detecting and classifying non-covalent interactions between a nucleic acid (RNA or DNA; called a *receptor*) and *ligands* (metal cations, small molecules, nucleic acids, or proteins). These data are encoded in the form of Structural Interaction Fingerprints (SIFts) - a 1D bit vector indicating presence or absence of a given type of interaction, as well as in the form of a detailed listing of all detected interactions, spatial coordinates of the interacting partners, and distances between interacting atoms or aromatic rings.

Here we present three analyses performed for RNA-ligand complexes. In all the cases, the fingeRNAt played a pivotal role in data gathering and analysis.

### Classification of interactions in experimentally solved RNA-ligand structures

Experimentally solved structures of macromolecules and their complexes are an invaluable source of knowledge on intermolecular interactions. At the time this publication was written, 1570 structures of RNA had been deposited in the Nucleic Acid Database, with 946 structures of RNA complexes with small molecule ligands (as of 16-Dec-2021, (45)). The information on statistics of interactions in RNA-ligand complexes derived from the solved structures can be used to develop bioinformatics methods to predict the structure of such complexes. Methods that enable an analysis of RNA-ligand interactions include docking programs (such as rDock (46,47)) or scoring functions (such as DrugScoreRNA (48), RNAPosers (49), and developed in our laboratory LigandRNA and AnnapuRNA (50,51)). As an output, the aforementioned methods return the proposed binding pose with numerical score(s). Although these data offer great help in compound prioritization processes or virtual screening, they do not explain the nature of the binding phenomena nor give insights into the main driving forces of the investigated interaction. To shed light on the landscape of interactions with small molecule ligands, we analyzed a diversified dataset of experimentally solved RNA structures deposited in the PDB. We determined the nature of formed non-covalent interactions, including frequency and distance distribution for each investigated contact type.

The performed analysis reveals that the most frequently occurring interactions are hydrogen bonds (5026) and lipophilic interactions (3582; see Fig 1A and S2 Table). Next, with an order of magnitude lower number, are cation-anion bonds (899), water-mediated interactions (151), and Pi-stacking interactions (146). The number of the remaining interactions is two orders of magnitude lower than the number of detected hydrogen bonds, with halogen bonds being the least frequent detected interaction (6). Three metal cations that were present only in one complex each (namely: Pb, Mn, and Sr) were removed from plots for clarity.

**Fig 1.**
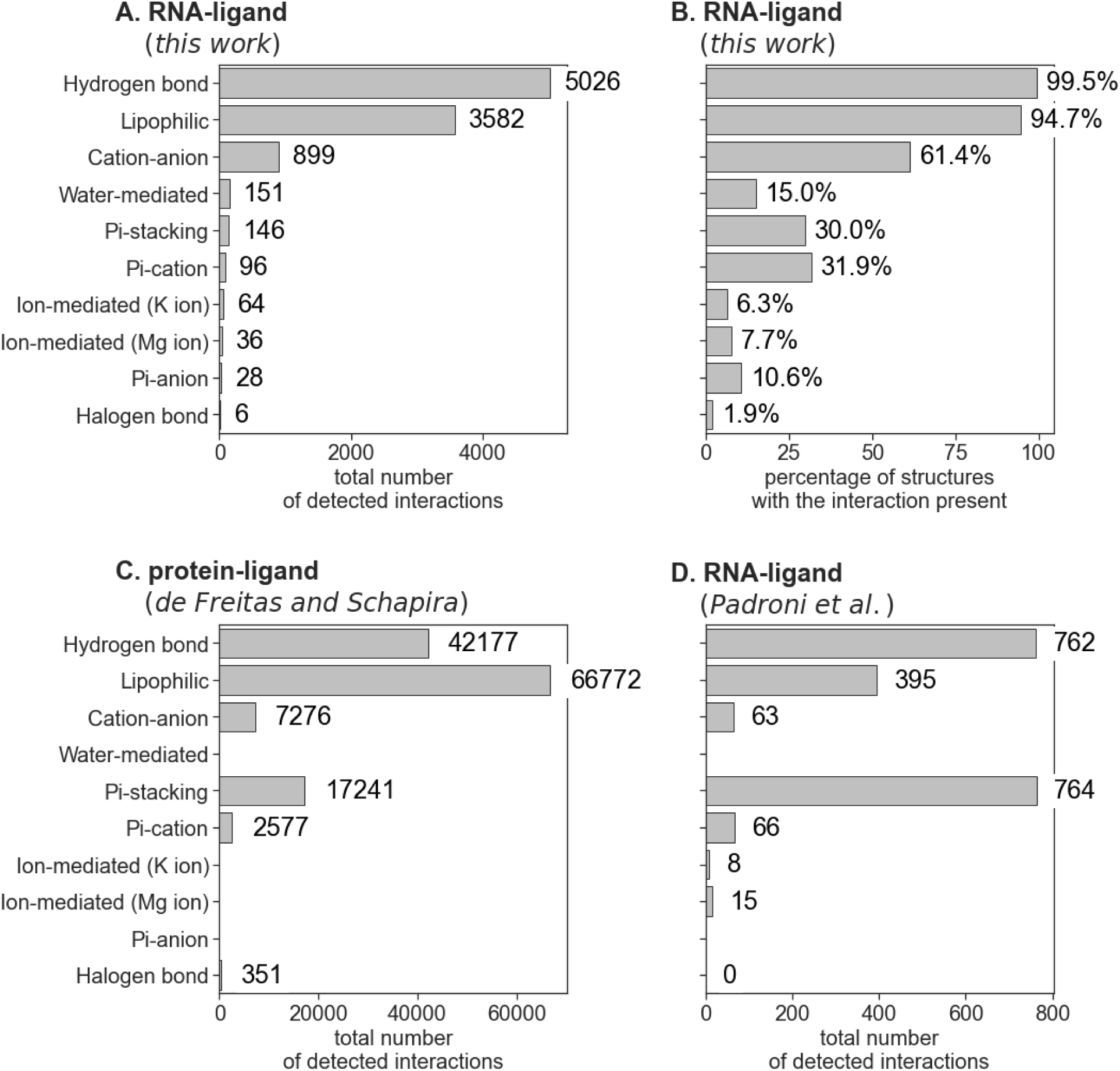
Statistics of interactions formed in macromolecular complexes. (**A**) Total number of interactions detected for RNA-ligand complexes (this work); (**B**) the percentage of RNA-ligand complexes with at least one occurrence of a given interaction (this work); (**C**) number of interactions for protein-ligand complexes presented by de Freitas and Schapira; (**D**) number of interactions for RNA-ligand complexes presented by Padroni *et al*.

Statistics indicating the absolute number of interactions, although informative, may be somehow misleading. Seven types of interactions (namely hydrogen bonds, lipophilic, cation-anion, halogen bonds, and mediated interactions) are detected between single atoms/groups of RNA and ligands. Thus, a single interaction point may form multiple bonds with the partner (e.g., a single lipophilic atom of RNA may form bonds with multiple atoms of the ligand and *vice versa*). Interactions involving Pi-systems (Pi-stacking and Pi-ions interactions) are formed between at least one cyclic system, which due to geometrical restraints may form only a limited number of interactions.

To circumvent this bias, we calculated the number and the percentage of complexes with at least one of the given interactions detected (see Fig 1B). This analysis revealed that the hydrogen bonds, lipophilic, and cation-anion interactions are still the most abundant, present in over 99%, 94%, and 61% of structures, respectively. In this analysis, the role of the interactions formed by Pi-systems is more pronounced - the Pi-stacking and Pi-cation interactions are formed in over 30% of structures, and Pi-anion interactions are detected in 11% of structures. The percentage of structures forming mediated interactions is lower than suggested by the number of interactions formed (15%, 8%, and 6% for structures with water-, Mg^2+^-, and K^+^-mediated interactions, respectively). We also counted the number of interactions formed by unique residues of RNA (i.e., only one interaction of a given type with a given residue is counted). Results confirm the order of the frequency of interactions revealed by our first analysis (Fig 1A), but with a slightly decreased role of water-mediated interactions (S1 Fig).

Comparison of the RNA-ligand interactions statistics to the data derived from proteinligand complexes published by de Freitas and Schapira reveals that the two most frequent interactions are the same (hydrogen bonds and lipophilic interactions), however in the reversed order (Fig 1C). To our surprise, the Pi-stacking interactions are more pronounced for protein complexes than for RNA. This observation may be explained by the fact that only 38% of ligands in the RNA-ligand dataset contain an aromatic ring, which is a prerequisite for forming the Pi-stacking interaction with RNA (S2 Fig; data for protein complexes are not published), and almost 80% of ligands with an aromatic ring is forming a Pi-stacking interaction with RNA (see S3 Table for detailed statistics of Pi-stacking interactions, S4 Table for hydrogen bonds, and S5 Table for cation-anion interactions).

We also compared statistics of the interactions derived using the fingeRNAt to data published recently by Padroni *et al*. (52). The authors used a proprietary software ICM to extract contact information from a set of 37 experimentally solved structures, which covered 14 unique RNAs (data summarized in Fig 1D). They found the Pi-stacking interaction and hydrogen bonds as the most common type of interaction in RNA-small molecule ligand complexes, with an almost equal number of detected interactions (764 and 762, respectively). Our analysis also ranks the hydrogen bonds as the most frequently occurring interaction, but as discussed above, the Pi-stacking interaction is less commonly discovered (the fifth most frequent interaction detected by our method). The third most frequent contact found by Padroni *et al*. is the lipophilic interaction (the second most frequent interaction detected by our method). The substantial difference in contact count values is, however, in the number of cation-anion interactions. In our analysis, this is the third most frequent interaction, while Padroni *et al*. rank it in the fifth position. We also detected four cases of halogen bonds, while Padroni *et al*. did not observe this kind of interaction. Observed discrepancies in absolute values and frequency ranks may result from the fact that the analysis presented by Padroni *et al*. was performed on a relatively limited and not diversified set of structures (37 structures vs. 207 diversified structures analyzed in this work), thus introducing a possible bias to the statistics of preferred interactions. What is more, we used protonated ligand molecules as an input which more realistically reflects the ionization state of the binding partners and their interactions, thus explaining the differences in the charge-involving interactions (cation-anion, Pi-cation interactions).

We also used the data generated by the fingeRNAt to investigate the preferred distances for detected interactions. The distribution is multimodal for hydrogen bonds and cation-anion interactions, with two clearly distinguished peaks (Fig 2, S3 Fig, and S6 Table). Such distribution was observed earlier for strong hydrogen bonds (for data derived from structures deposited in the Cambridge Structural Database (CSD) (53,54) and protein-ligand complexes (55)), metal cations with oxygen anion (56), salt bridges in proteins (57), and Pi-anion interactions (58). For hydrogen bonds length distribution, the first peak is observed for the distance 2.7 Å, which is very close to the median length observed for hydrogen bonds in data derived from the PDB and CSD (2.75 Å and 2.9 Å for interactions of amide C=O with OH and NH, respectively, (59)). We observed the second-main peak at the length 3.7 Å, which may result from water-mediated contacts via water molecules not visible in the experimentally determined structures.

**Fig 2.**
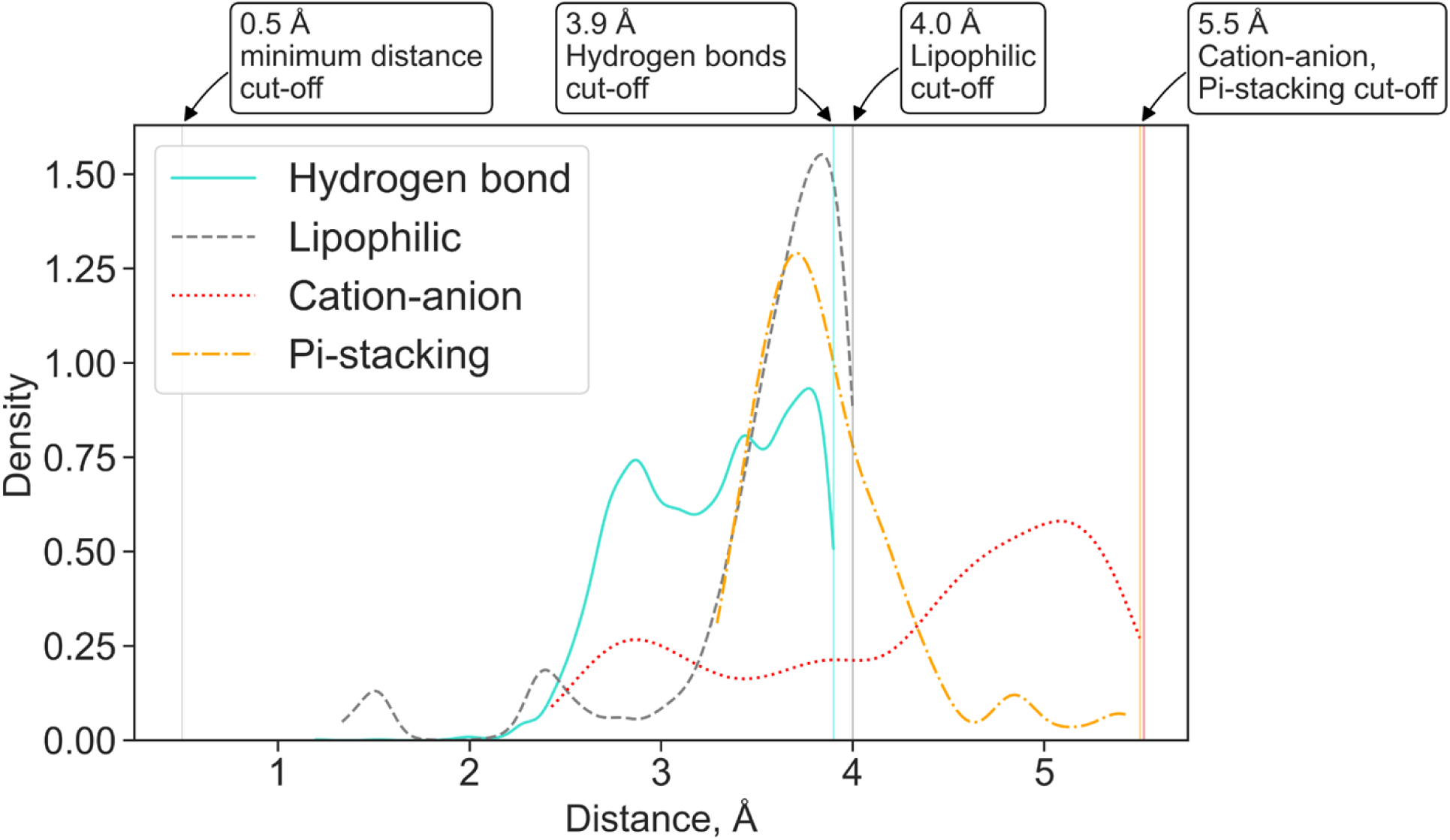
A kernel density estimate plot of the distribution of lengths for four most frequently observed non-covalent interactions in a dataset of experimentally solved RNA-ligand structures. For hydrogen bonds, the distance between the non-hydrogen donor atom and the acceptor is reported. The minimum and maximum cut-off values for each interaction are marked with vertical lines.

Using the interaction statistics gathered with the fingeRNAt software, we calculated the frequency of interactions formed by the individual RNA atoms. We also mapped the data into the nucleotide structures to visualize the Interactions’ Hot Spots. For hydrogen bonds, most of the hydrogen bonds are formed by nucleobases (61%) and phosphate group atoms (23%), while ribose oxygen atoms are responsible only for a small fraction of the hydrogen bonds (15%, see Fig 3 and S7 Table). This observation is in agreement with results obtained by Kligun and Mandel-Gutfreund, indicating that nucleobases form 65%, while interactions with backbone atoms form 35% of RNA-ligand hydrogen bonds (60). Our observation confirms that the majority of RNA-ligand hydrogen bonds are nucleobase-specific.

**Fig 3.**
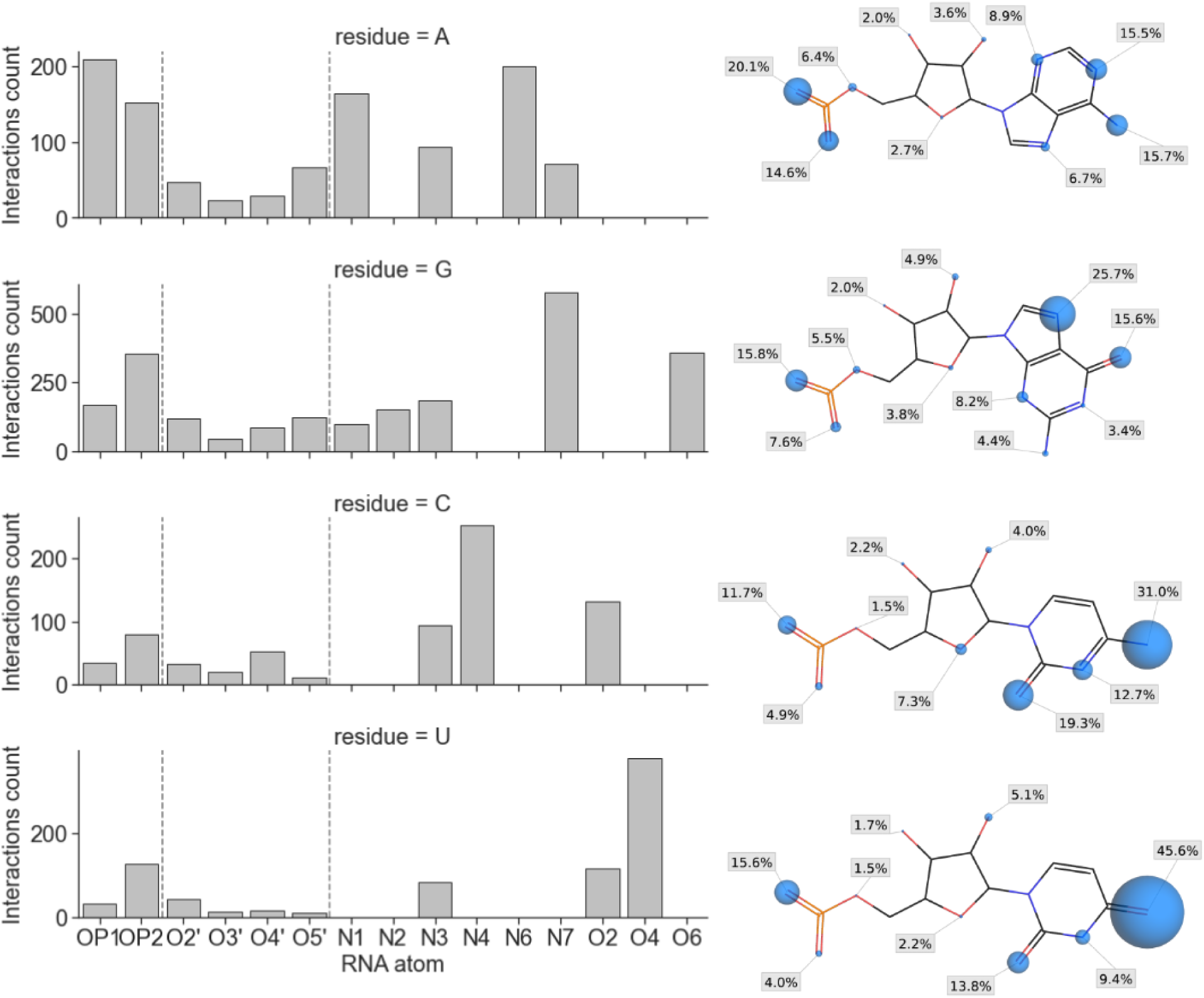
The number of hydrogen bonds formed with ligand molecules grouped by the RNA atoms (bar plots, left; vertical dashed lines separate atoms of a phosphate group, ribose, and nucleobase) and interaction sites statistics for each atom with a percentage of all hydrogen bonds formed by this residue (right column; the radius of spheres are proportional to the percentage values).

Analysis performed for lipophilic interactions indicates that most contacts between RNA and ligands are also made with nucleobase atoms (74%, Fig 4 and S8 Table). For Pi-anion interactions, 82% of bonds are formed with nucleobases, with RNA acting as an anion acceptor (in the remaining 18% of Pi-anion interactions, the phosphate group of RNA acts as an anion donor; see S9 Table). Interestingly, we observed the formation of a halogen bond mostly with the ribose atoms (5 cases, 83%) with a single interaction detected with a nucleobase atom (see S10 Table). However, the overall number of these interactions in our dataset is low (6 interactions), therefore the observed trend may not be reliable. Due to the molecular features of the RNA, all Pi-stacking and Pi-cation bonds are formed exclusively with nucleobases. Taken together, most of the observed interactions of all kinds are formed with the nucleobases (60.53%, while 21.63% and 17.84% with phosphate and ribose fragments, respectively), which highlights a vital role of this region of RNA in structure recognition (see S11 Table).

**Fig 4.**
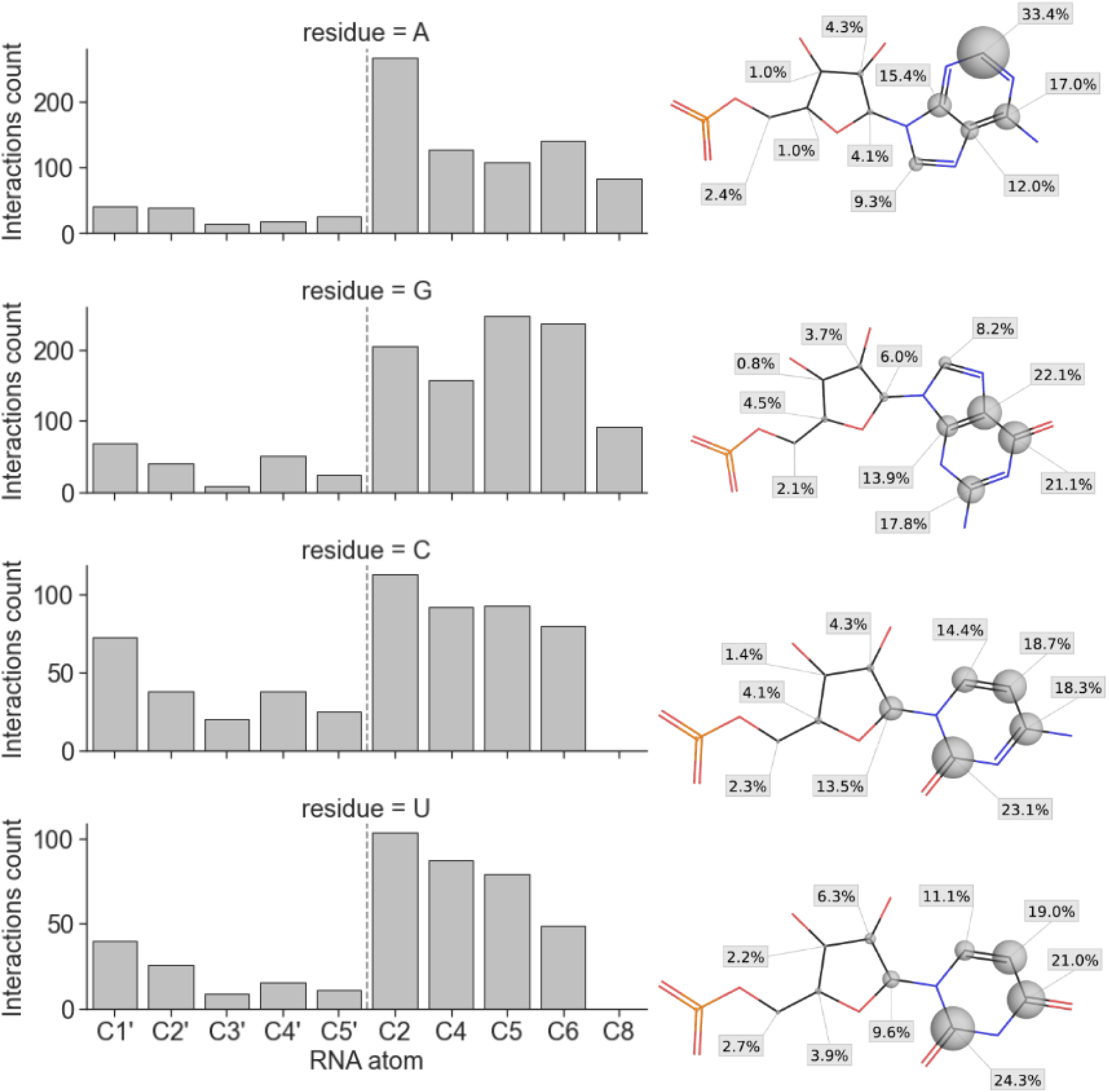
The number of lipophilic interactions formed with ligand molecules grouped by the RNA atoms (bar plots, left; vertical dashed lines separate atoms of ribose and nucleobase) and interaction sites statistics for each atom with a percentage of all hydrogen bonds formed by this residue (right column; the radius of spheres are proportional to the percentage values).

Presented data can pave the path toward a better understanding of the nature of such interactions and define the main driving forces responsible for forming these complexes. Medicinal chemists can directly use provided information on the preferred interaction distances and interaction sites to support the rational design or modification of existing small molecule ligands to improve their binding affinity or selectivity toward RNA molecules.

The absolute number of detected interactions, however, must be treated as an approximation only. As recently shown by Xu *et al*. for protein-ligand and protein-protein complexes, insufficient resolution of the data deposited in the PDB may result in overlooking a significant number of non-covalent interactions (61). Most probably, this problem also exists for RNA-ligand complexes; still, the presented data gives an overview of the relative abundance of the non-covalent bonds in examined structural data.

### SIFt-based structure similarity assessment

The most widely used criterion of the accuracy of macromolecule-ligand modeling tools is the ability to reproduce the binding mode of ligands. This is usually measured by calculating the root-mean-square distance (RMSD) between the non-hydrogen atoms of the ligand in the experimentally determined structure and the corresponding atoms in the modeled pose. Although widely used for the assessment of molecular docking programs (62), where the macromolecule structure is usually kept rigid, RMSD has several shortcomings, mostly seen in simulations involving the flexibility of both interacting partners. It may happen that although the structure of the predicted complex is very similar to the reference structure and the binding mode of the ligand is perfectly recapitulated, the RMSD value is very high (also poor). In addition, the RMSD is ligand-size dependent, tangling direct comparison of RMSD values obtained for molecules of different sizes as well as using a fixed RMSD threshold as a criterion for successful molecular docking.

To circumvent the above mentioned drawbacks of RMSD, several alternatives have been proposed, however, they were designed and tested, to the best of our knowledge, exclusively for protein complexes. The list includes methods such as RSR (Real Space R-factor, which measures how well a group of ligand atoms fits the experimental electron density, (63)), GRAD (Generally Applicable Replacement for rmsD, which takes into account relative importance to binding of atoms, (64)), TFD (Torsion Fingerprint Deviation, which compares conformations of molecules (65)), or SuCOS (for assessing shape complementarity and overlapping of chemical features (66)). Also, several metrics utilizing a comparison of contacts between proteins and ligands have been proposed. Ding *et al*. developed the Contact Mode Score (CMS), a metric to assess the conformational similarity based on intermolecular protein-ligand contacts (67). CMS is expressed as the Matthews correlation coefficient (MCC) between contact matrices generated for the reference structure and the investigated complex. It was shown to be a valuable metric to evaluate results of flexible docking, which at the same time considers the changes upon ligand binding. In the IBAC approach (Interactions-Based Accuracy Classification, (68)) proposed by Kroemer *et al*., the scoring is derived from the comparison of (manually defined) key interactions of the reference protein-ligand complex and the docked pose. Although defining the key interactions may be perceived as subjective, the IBAC method was proved to be a more meaningful measure of docking accuracy for the examined test set than RMSD. Balius *et al*. proposed an FPS score (footprint similarity, (69)) which is derived from the comparison of electrostatic, steric, and hydrogen bonding energy profiles for protein-ligand complexes using the Pearson correlation coefficient.

The similarity of interaction fingerprints for protein-ligand complexes was also explored as a measure for binding mode similarity. Drwal *et al*. analyzed four protein targets to compare binding modes of fragments, crystallization additives, and drug-like molecules (70). For a given target, they calculated a consensus fingerprint containing the relative frequency of each interaction type with each residue. The similarity between a docking pose fingerprint and a consensus fingerprint was calculated using the Tanimoto metric for continuous variables. Leung *et al*. benchmarked the PLIF similarity (Protein-Ligand Interaction Fingerprint) as a metric for evaluating docking of the ligands to proteins (66). They concluded that this metric, contrary to ligand-centric ones (such as the RMSD and SuCOS), was able to capture information about interactions across multiple crystal structures of ligands bound to the same protein, making it a handy feature for experiments where multiple protein conformations are used.

Inspired by structure-centric methods developed for assessing protein-ligand complexes, we examined the applicability of the interaction fingerprints generated by the fingeRNAt as a measure for RNA-ligand complexes’ similarity. As input structural data, we used predictions submitted by participants of the RNA-Puzzles - a collective experiment for blind RNA structure prediction. In the RNA-Puzzles round 23, the task was to predict structures of a Mango-III aptamer in complex with biotinylated TO1 dye, based on the sequence of RNA and structure of the ligand. The main criterion of the evaluation of the prediction’s quality was the RMSD, deformation index (DI), and interaction network fidelity (INF, (71)) of RNA. Seven groups submitted their models of the target complex (RNA with ligand) - Bujnicki, Chen, Das, Ding, Dokholyan, Adamiak (codenamed RNAComposer), and Xiao. The structures of ligands provided by the last group contained structural errors, which made it impossible to calculate RMSD values for the ligand; however, the calculation of interaction fingerprints was possible due to the structure-agnostic (structure-independent) nature of the algorithm proposed in this work.

First, we compared the correlation of the RMSD and INF of RNA and RMSD of the ligand with interaction fingerprint similarity. The similarity of interaction fingerprints (and, in general, of bit vectors) can be expressed in a number of ways. Currently, the fingeRNAt package offers eight methods for calculating the similarity or distance of bit vectors. Here, we used a widely used metric for the comparison of interaction fingerprints - the Tversky similarity (66,72). It focuses on the recapitulation of the true interactions (formed in the reference complex) in the predicted model of a complex and ranges from 0 (investigated model has no interactions which are present in the reference complex) to 1 (model has all interactions the reference has).

We observed only a weak correlation between RMSD (of RNA or ligand), INF, and Tversky similarity (R^2^ for linear least-squares regression ranges from 0.31 to 0.61; see Fig 5A and S12 Table). The detailed analysis of the rank list shows that most of the best models in terms of RMSD, DI, or INF do not recapitulate the interaction network formed between RNA and ligand in the experimentally solved structure (S13 Table). Only the model submitted by the Das group (Das_07) had relatively good values of RMSD (for both RNA and ligand), DI, and interaction fingerprint similarities (S4 Fig). Also, the mutual similarity of interaction networks in the submitted models is relatively low (with average Tversky similarity equal 0.24 and median similarity 0.18), indicating that in most complexes, the proposed binding mode is unique (S5 Fig).

**Fig 5.**
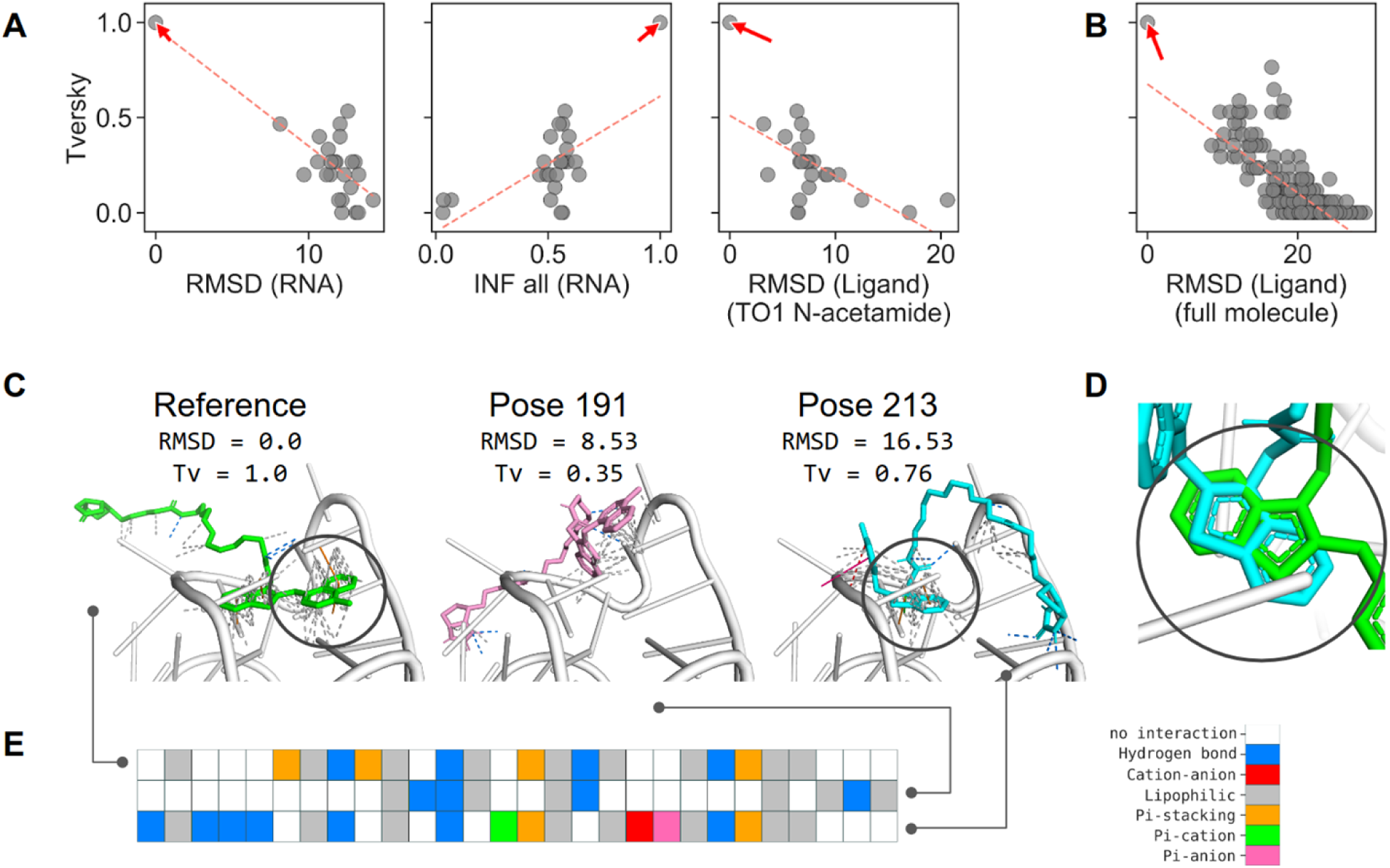
Relationship between structure-centric similarity metrics (RMSD and INF) and fingeRNAt interaction fingerprints similarity (Tversky). (**A**) Relationship between RMSD of RNA, INF all for RNA, RMSD of ligands, and Tversky similarity of interaction fingerprints for models submitted for the RNA-Puzzles round 23 and (**B**) for the redocking experiment. The reference structure is marked with an arrow, and the linear least-squares regression is marked with a red dashed line. (**C**) Selected poses from the docking experiment with ligand RMSD and Tversky similarity of fingerprints, (**D**) the overlay of the benzothiazole fragment of the ligand of the reference structure, and pose 213, and (**E**) the corresponding color-coded interaction fingerprint. Benzothiazole moiety was marked with a gray circle in (C) and (D).

As a complementary experiment, we performed molecular redocking of the ligand to the reference structure of the Mango-III aptamer. Again, we observed only a weak correlation between RMSD of ligand and Tversky similarity of interaction fingerprints (R^2^ for linear leastsquares regression equal 0.60; see Fig 5B and S12 Table). For the detailed analysis, we selected two poses proposed by the docking program - one with relatively low (good) RMSD of ligand but low (unfavorable) fingerprint similarity (pose 191) and one with relatively high (unfavorable) RMSD value but high (good) fingerprint similarity (pose 213) (Fig 5C). Comparison of the two selected poses with the reference complex confirms that pose 191, despite its better RMSD value, has a very different binding mode than the reference ligand. In this case, only six out of 17 interactions were correctly predicted. Conversely, pose 213 with a very high RMSD value recapitulates 13 out of 17 interactions formed by the reference structure. This is especially pronounced by the correct prediction of the position of the central benzothiazole ring, although in a different orientation (Fig 5D).

Moreover, for the redocking data, we did not observe the strong correlation between RMSD of the ligand and any of the eight metrics expressing similarity/distance of interaction fingerprints (for the mutual relationship plots of RMSD and bit vector similarity/distance values, see S6 Fig), meaning that the assessment method proposed by us is distinct from other metrics and could be easily employed as another mean of structure’s prediction scoring. Also, lower resolution fingerprints (SIMPLE, PBS) can be used for the rough estimation of the binding mode - the similarity of these fingerprints does not correlate with the RMSD of the ligand as well (S7 Fig).

In summary, the similarity of the interaction fingerprints can be used as an alternative to the RMSD measure for comparing structures of complexes. The proposed approach focuses on the interactions formed between interacting partners, while the actual location of particular atoms and groups of ligands is not considered. By using different fingerprint similarity measures, various aspects of interaction similarity can be emphasized. While the Tversky similarity expresses the number of correctly predicted interactions, the Tanimoto similarity expresses the overall mutual similarity of the interactions in compared complexes.

These results are in line with data obtained earlier for protein-ligand complexes. It was shown that the interaction data could improve docking accuracy by selecting the binding pose closer to the reference structure and recapitulate models poorly assessed by other methods (73,74).

### Fingerprints for clustering interactions and detecting preferred patterns

As stated in the previous section, RMSD is the most widely used measure for assessing the similarity of ligands in complex with macromolecules. It may be treated as a distant proxy for comparing interactions formed in two investigated complexes, assuming that two molecules that are close in space will form the same (or similar) interaction network. As we showed, this is not always true, as two ligands with low values of RMSD may form very different interactions with the receptor and *vice versa*. Another limitation of the RMSD used for comparing the binding modes in two complexes is the restriction that both ligands must be the same. Although some recently published methods partially circumvent this constraint by enabling the calculation of the RMSD of molecules sharing some structural features (LigRMSD, (75)), application of RMSD is still limited to structurally similar compounds.

In this experiment, we tested the applicability of SIFts for grouping small molecule compounds of different chemical structures but with akin binding patterns. We hypothesized that molecules forming similar types of interactions with the molecular target would have similar binding properties, such as biological activity. We composed a dataset of diversified small molecule ligands with known or putative activity toward the HIV-1 trans-activation response (TAR) element. The library consisted of 30 active and 1478 inactive molecules (see Materials and methods section Datasets for the detailed description of the library preparation). Using molecular docking, we predicted the structure of these compounds in complexes with HIV-1 TAR RNA and calculated SIFts for each RNA-ligand complex.

In a pool of 1508 docked compounds, there were 1149 unique fingerprints, and 998 compounds (66.2%) had a unique fingerprint (i.e., the fingerprint that is unique for this molecule only). All 30 active compounds had a unique fingerprint. We used PCA for dimensionality reduction and *k*-means clustering for grouping compounds with similar fingerprints, i.e., forming a similar interaction network with the target RNA. This method offers a reasonable separation of clusters (average silhouette score of 0.516 for 15 clusters, Fig 6A and 6B; for the performance of clustering for various numbers of clusters, see S14 Table). In five clusters, the ratio of active compounds was significantly different than in the input dataset (*p*-value ≤ 0.05, clusters 8-12), although it did not include the cluster with the highest ratio of active compounds (cluster 7, 6.67% of active compounds, *p*-value = 0.29, Fig 6B; for clusters composition and statistical significance analysis, see S15 Table). In clusters 8, 11, and 12, the ratio of active compounds is significantly higher than in the input population (with the percentage of active compounds 2.38%, 4.51%, and 4.00%, respectively, *p*-value ≤ 0.05). Thus, we conclude that the interaction pattern present in the latter three clusters may be “favorable” for active compounds. Conversely, two clusters (9 and 10) do not contain any active compounds (and this number is significantly lower than in the input population), which could indicate that interaction networks formed by members of this group are “unfavorable” for HIV-1 TAR binding ligands.

**Fig 6.**
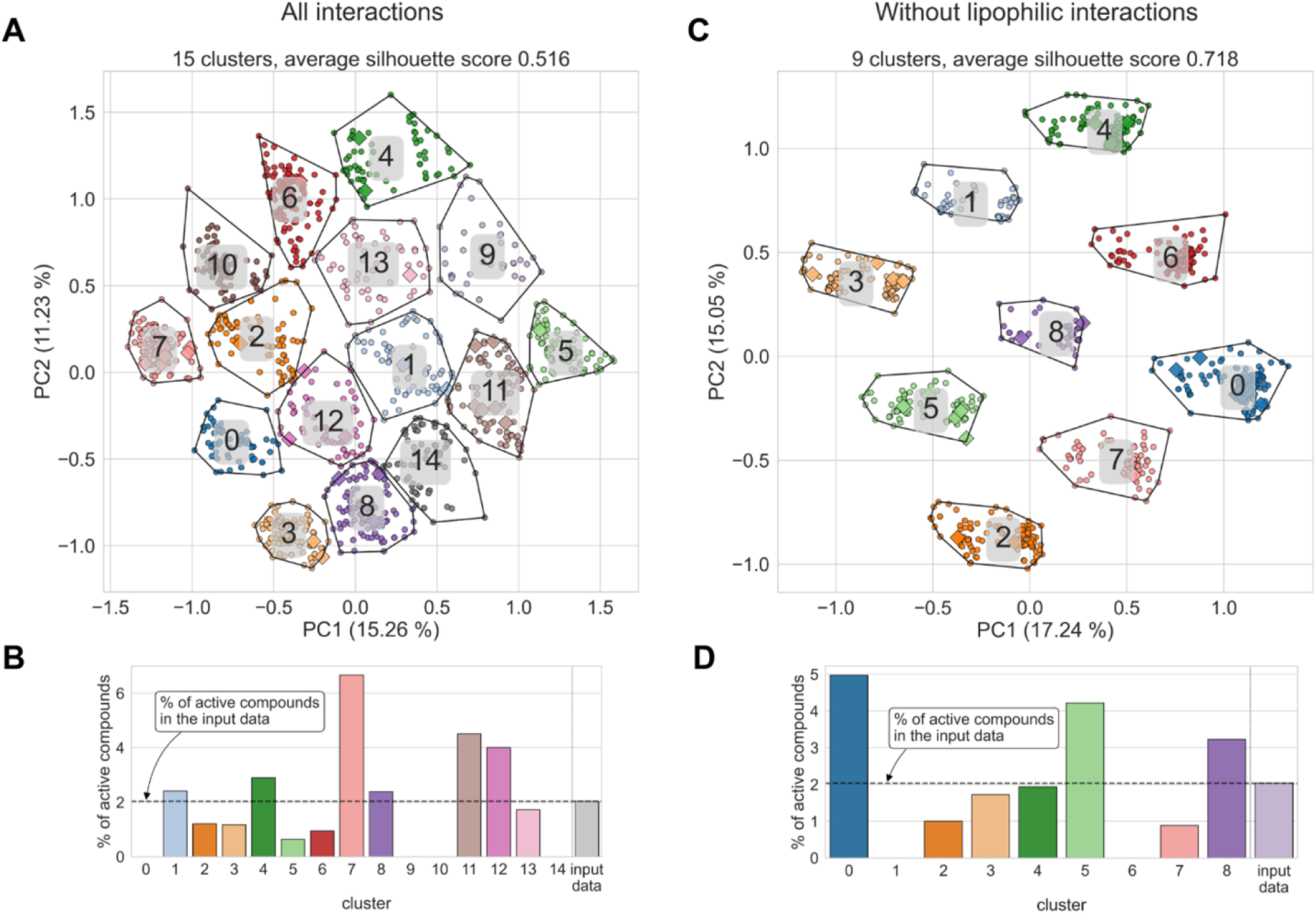
Interaction fingerprints used for visualization and clustering of predicted interaction patterns calculated for data from molecular docking of a set of active and inactive compounds to HIV-1 TAR structure (1UTS). (**A**) Interaction fingerprints mapped on the 2D space using PCA with color-coded clusters for the all-interactions dataset and (**C**) with lipophilic interactions removed. (**B**) Percentage of active compounds in each detected cluster compared to the value obtained from the input dataset, calculated for all-interactions dataset and (**D**) obtained from the dataset with lipophilic interactions removed.

We performed the same type of analysis for the dataset with lipophilic interactions removed. These interactions are most common among all complexes, and we hypothesized that they might introduce noise into the fingerprint data. If the lipophilic interactions are removed, the separation of the clusters is even better (average silhouette score of 0.72 for nine clusters, Fig 6C and 6D). We detected two clusters in which the ratio of active compounds was significantly lower than in the input dataset (*p*-value ≤ 0.05, clusters 1 and 6) and which did not contain any active ligand. The ratio of active compounds in cluster 0 was 4.97%, which is 2.5 times higher than in the input dataset (however, this result is not statistically significant, *p*-value = 0.08). This analysis enabled us to define nine groups of interaction patterns formed by the ligands in complex with HIV-1 TAR, some of which are more “favorable” or “unfavorable” for active ligands. Comparing the results of this analysis with the previous one, where all interaction types were present, we conclude that lipophilic interactions introduced noise to the fingerprint data, making generated clusters fuzzier. On the other hand, it enabled a better separation (more distinctive clusters) of active and inactive compounds.

Taken together, dimensionality reduction and clustering of fingerprints offer a high-throughput method of analysis of structural data of complexes. As shown, it may be used to define groups of ligands interacting with a receptor in a similar way and thus sharing similar properties (such as biological activity or lack of thereof). This kind of analysis may also be used in high-throughput virtual screening to generate a diversified set of compounds (in terms of forming diverse interactions with the receptor of interest) for further testing for their biological activity. Clustering fingerprint data may help select groups of compounds forming similar interactions to the active compounds, thus having a higher probability of being biologically active. It could be directly applied as a post-processing step of virtual screening with molecular docking. Decomposition methods other than PCA may be used, leading to potentially meaningful results (such as obtaining clusters enriched with molecules with a given property; see S8 Fig). Contrary to the ligand-based metrics (such as the RMSD), the presented approach is not limited to the same or structurally similar ligands.

## Summary

Interactions between nucleic acids and ligands play a pivotal role in many biological processes. Characterization of these interactions may elucidate our understanding of those phenomena and help to explain the nature of molecular recognition. This knowledge may also be utilized to modulate the binding process in the desired way, for example, by small molecule inhibitors binding to RNA and preventing its interactions with a molecular partner. The presented software tool, fingeRNAt, detects and characterizes interactions in nucleic acid-ligand complexes. We showed its applications in different bioinformatics problems to help answer structural biology and drug development questions. These included analyzing experimentally solved RNA-small molecule ligand complexes deposited in the PDB database to determine the statistics of non-covalent bond types and their features. We also proposed SIFts’ similarity as an alternative measure to RMSD. Contrary to the RMSD, SIFt-based metrics do not depend on the receptor conformation nor the ligand structure and focus on how well the interaction network is recapitulated in the model compared to the reference complex. Besides, we presented an application of molecular fingerprints for the clustering of complexes. Fingerprint data, processed with multidimensional scaling and clustering, yields groups of complexes with similar binding patterns. We demonstrated that these clusters might be enriched with compounds with desired properties, such as biological activity, facilitating a high-throughput analysis of the structure-activity relationship and visual analyses of multiple complexes. The accompanying PyMOL plugin enables visualization of the detected interactions, an inspection of the results, and preparation of publication-quality images.

fingeRNAt is relatively fast. Calculation of fingerprint type FULL for the redocking of guanidine ligand to guanidine III riboswitch (RNA with 39 residues, ligand with 10 atoms, 100 ligand poses) took less than 7 seconds, while calculating the Tanimoto similarity matrix took under 2 seconds (calculated on Ubuntu Linux 20.04 with Intel(R) Core(TM) i5-8400 CPU and 32 GB RAM; see S9 Fig and S16 Table for the detailed benchmark).

Applications of the fingeRNAt-generated SIFts significantly go beyond the ones described in this manuscript. The program may be used to generate interaction profiles of nucleic acid with ions, which may help understand ion binding preferences and enable comparing ion profiles between multiple structures. Moreover, fingeRNAt ideally fits the pipeline for analysis of molecular dynamics trajectories, indicating forming and breaking non-covalent bonds during a simulation. SIFts, paired with the bioactivity data for small molecules, may also be used to develop predictive models for the molecular target of interest.

The roadmap of the software development includes detection of less frequently observed types of non-covalent bonds, such as halogen-Pi (76), “mixed” type bonds, such as cation-anion hydrogen bonds (77), or a separate class of anions binding to amino, imino, and hydroxyl groups of RNA (78). Recent publications suggest that these interactions may greatly contribute to the molecular recognition process (for a recent review on “unusual” interactions, see (79)). New levels of the resolution of the fingerprint will include differentiation of strong and weak interactions (e.g., for hydrogen bonds). We also plan to define geometrical rules for ions and water molecules to more reliably detect such interactions.

We are confident that fingeRNAt will be highly useful for the bioinformatics community and will facilitate research on nucleic acid interactions.

## MATERIALS AND METHOD

### The fingeRNAt method

fingeRNAt is a set of Python 3 programs for detection, classification, and analysis of interactions formed within nucleic acid-ligand interactions. It consists of three main tools, each serving a different purpose.

#### fingeRNAt.py

fingeRNAt.py is a program for the detection and classification of non-covalent nucleic acid-ligand interactions. It can be run from the command line or via the graphical user interface. As an input, it takes a 3D structure of a receptor (RNA or DNA) and a file containing ligands, which form a complex with this macromolecule. fingeRNAt.py accepts six types of ligand molecules: small molecules, proteins, metal cations, DNAs, RNAs, and LNAs (locked nucleic acids) (Fig 7). The output is a fingerprint - a bit vector containing information on the declared interactions detected between the receptor and the ligand.

**Fig 7.**
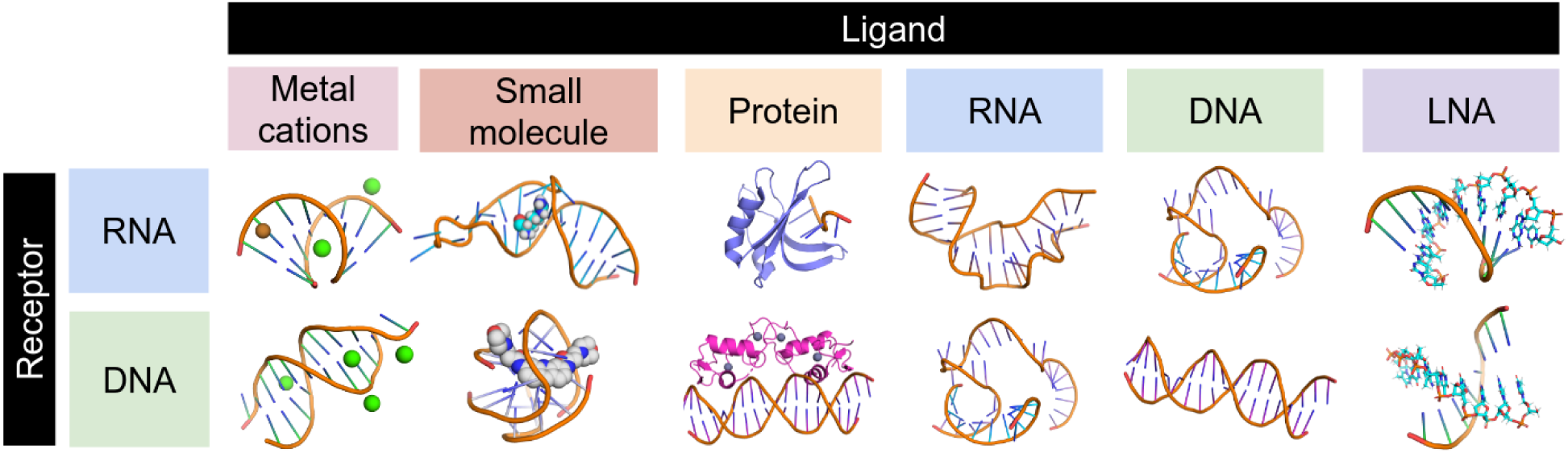
Combinations of receptor and ligand types accepted by the fingeRNAt.py.

#### Input

For all ligand types but metal cations, the program requires two input files: (i) a receptor file, which is an RNA or DNA structure in the pdb format (one model per file), with explicit hydrogens added, and (ii) a ligand file in sdf format, which may contain multiple structures. It is possible to calculate profiles of inorganic ions’ interactions with the receptor; in such a case, only one input file containing an RNA or DNA structure in the pdb format should be passed, and inorganic ions should be present within the same file. fingeRNAt will then treat all inorganic ions as ligands and calculate SIFt for each residue - ion pair.

The receptor’s structure may contain water and metal cations, but any ligands or buffer molecules must be removed prior to the analysis. Input ligand molecules must have assigned desired protonation states and formal charges. Formal charges on the phosphate groups do not need to be explicitly indicated in the receptor molecule, as fingeRNAt.py always treats OP1 and OP2 atoms as negatively charged anions.

#### Output

The output is a SIFt calculated for each nucleic acid-ligand pose, saved in a tab-separated (tsv) file, with separate columns for each residue and interaction type. Optionally, the human-readable file (also in tsv format) with detailed information on detected interactions can also be created (-detail option), which includes a listing of all detected interactions, spatial coordinates of the interacting partners, and distances between interacting atoms or aromatic rings.

#### Implementation

For each ligand in the input file, the program iterates over all nucleic acid’s residues and detects interactions of a given type. If the interaction is detected, the respective bit in the fingerprint is set to “1” and to “0” otherwise. Interactions can be detected at three resolutions: (i) low-resolution SIMPLE variant detects contacts between any atom of each of nucleic acid residue and ligand; (ii) medium-resolution PBS variant detects interactions between atoms of Phosphate, Base, and Sugar fragments of a nucleic acid residue and a ligand; (iii) high-resolution FULL variant detects and classifies the type of non-covalent interactions between a nucleic acid and a ligand; in this variant, fingeRNAt detects nine hard-coded interaction types: (i) hydrogen bonds, (i) halogen bonds, (i) cation-anion interactions, (iv) Pi-cation interactions, (v) Pi-anion interactions, (vi) Pi-stacking interactions, (vii) metal cation-mediated: magnesium, potassium, sodium, and other metal cation-mediated, (viii) water-mediated interactions, and (ix) lipophilic interactions. We distinguished magnesium, potassium, and sodium cations as those are the most prevalent metal cations in nucleic acid complexes (80,81); other metal cation-mediated interactions refer to interactions mediated by not the aforementioned ions.

Additional interaction types can be defined by plugins encoded in a human-readable yaml file. Interacting atoms or groups are defined by SMARTS patterns, separately for the receptor and ligand. Interaction criteria are defined based on the distance, distance and angle, or distance and dihedral angles between atoms/groups. The sample plugin file contains a definition of five interactions: (x) any interaction (any contact between nucleic acid and ligand), (xi) polar interactions, i.e., hydrogen bonds without angle restraints, (xii) weak polar interactions, i.e., weak hydrogen bonds without angle restraints, (xiii) n→π* interactions, (xiv) weak hydrogen bonds, and (xv) halogen multipolar interactions. Taken together, with the default configuration of the fingeRNAt.py, it is possible to detect 15 different interaction types.

For the summary of fingerprints’ variants and features, see Fig 8. Geometrical criteria for interactions were taken from the literature: ((82,83) for hydrogen bonds, (84) for halogen bonds, (85) for cation-anion interactions, (86,87) for interactions with p-orbitals: Pi-stacking and Pi-ion interactions, (88) for ion-mediated interactions, (89) for water-mediated interactions, (52) for lipophilic interactions), (90) for n→π* interactions, and the PLIP algorithm (protein-ligand interaction profiler, (28)). See S17 Table for the geometrical criteria summary.

**Fig 8.**
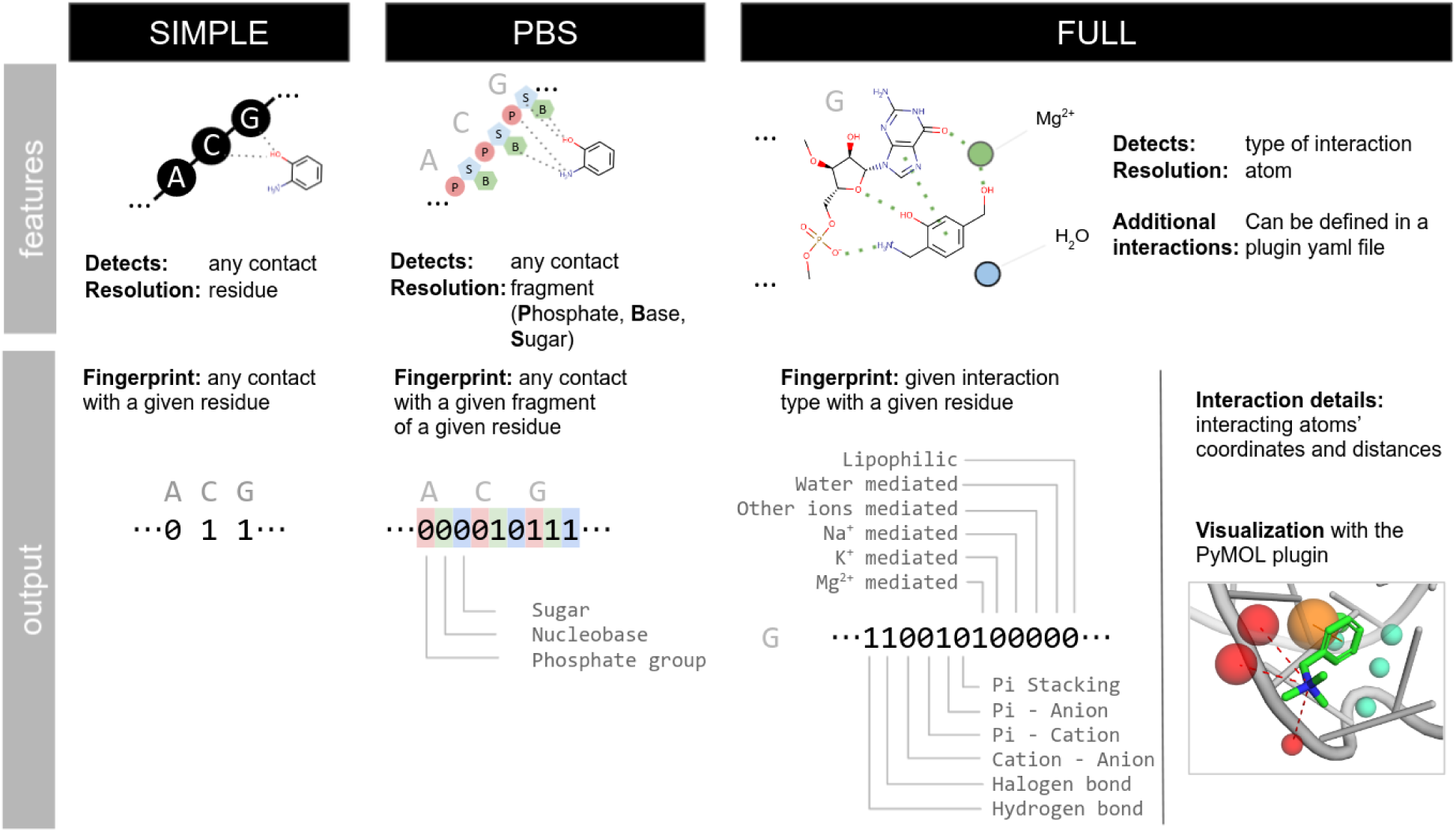
Fingerprints’ variants available in the fingeRNAt.py and their output.

Calculated fingerprints can be further post-processed by three wrappers: (i) ACUG for reporting interactions for four nucleotide types of the nucleic acid (A, G, C, and U or T), (ii) PuPy for reporting interactions for purines and pyrimidines, and (iii) Counter for reporting the total number of occurrence of a given interaction type. The detailed description of parameters accepted by the fingeRNAt is available in S18 Table.

fingeRNAt.py uses the OpenBabel Python module to parse input files and perform most cheminformatics calculations (such as hydrogen bonds acceptors/donors detection) and the RDKit Python module to detect aromatic rings and lipophilic atoms in ligands (91–93). A detailed description of the algorithm of fingerprints’ calculations is available in the program manual in the code repository. The summary of detected molecular features together with methods used can be found in S19 Table. If desired, the interaction definitions (such as distance threshold can be easily modified in a configuration file (for default values used in the program, see S17 Table).

#### fingerDISt.py

In addition to the main program, we provide an auxiliary tool, fingerDISt.py, to calculate different types of distances between fingerprints. It supports eight metrics: Tanimoto Coefficient, Cosine, Manhattan, Euclidean, Square Euclidean, Half Square Euclidean, Soergel (94), and Tversky distances (72). fingerDISt.py accepts SIFts’ tsv files generated by the fingeRNAt.py and returns a tsv file with a distance matrix for the fingerprints (all vs. all).

#### PyMOL plugin

The PyMOL plugin offers a convenient method of visualization of interactions detected and classified by the fingeRNAt program. After loading and processing the interaction file, the plugin generates three groups of objects: (i) Interactions, with objects representing detected interactions Fig 9A); (ii) Receptor Preferences (aka Receptor’s Interactions Hot Spots), with objects representing the spatial occurrence frequency of a given interaction type in receptor atoms (Fig 9B); (iii) Ligand Preferences (aka Ligand’s Interactions Hot Spots), with objects representing the spatial occurrence frequency of a given interaction type in ligand binding site (Fig 9C). Occurrence frequency is represented by spheres with centers at the interacting atoms, with the radius proportional to the ratio of interactions of a given type formed by the given interaction site to the total number of all interactions. Additionally, an auxiliary object representing only interacting residues of nucleic acid is created as well as a legend describing the color code and line style used for visualization of different types of interactions and Interactions Hot Spots (Fig 9D).

**Fig 9.**
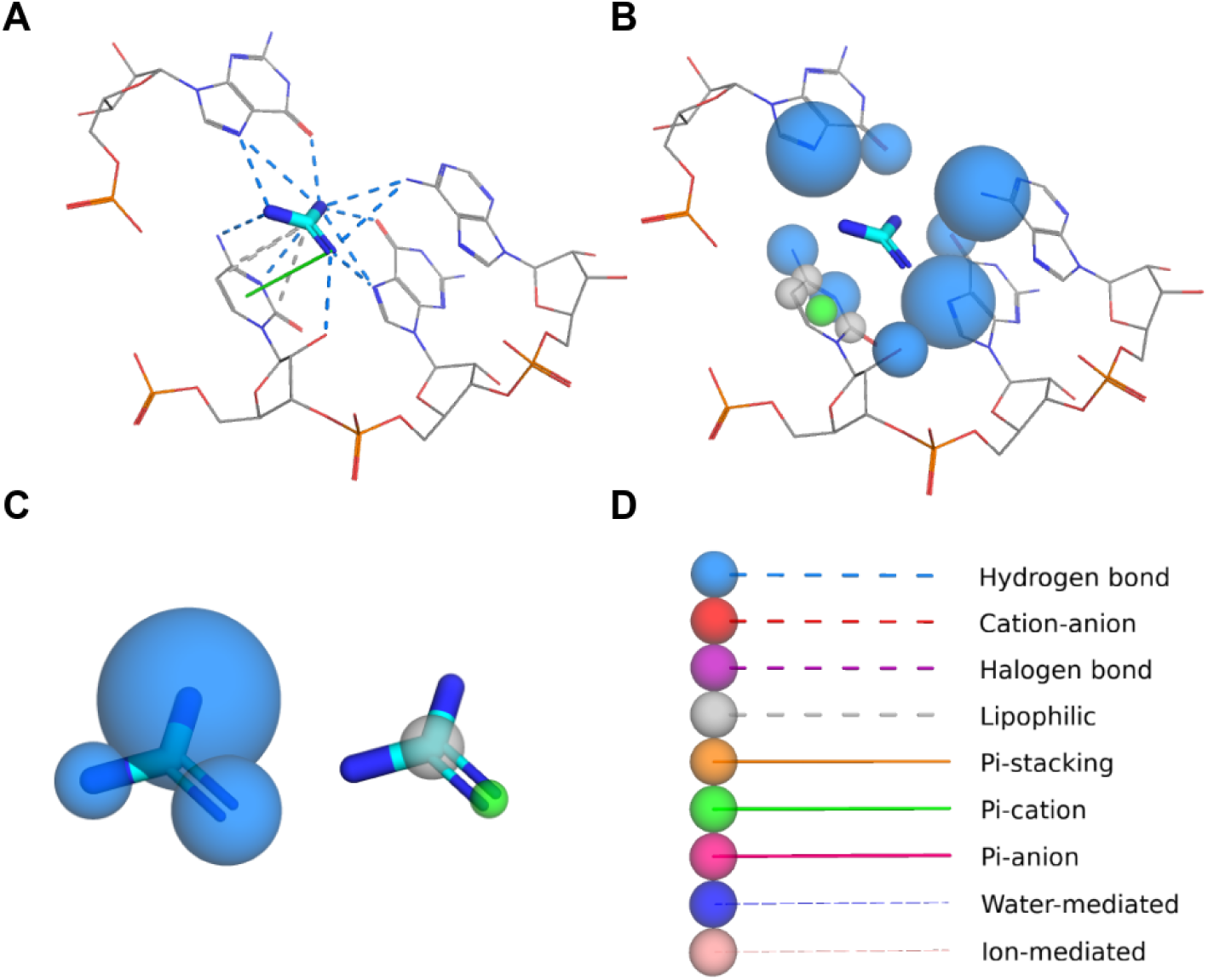
Three types of visualization generated by the PyMOL plugin. (**A**) Detected non-covalent interactions between nucleic acid and ligand; (**B**) Receptor Preferences (aka Receptor’s Interactions Hot Spots), with objects representing the spatial occurrence frequency of a given interaction type in receptor atom; (**C**) Ligand Preferences (aka Ligand’s Interactions Hot Spots), with objects representing the spatial occurrence frequency of a given interaction type in ligand atoms; (**D**) a legend describing the color code and line styles used for visualization of different types of interactions and Interactions Hot Spots.

#### Known limitations

The analysis of the X-ray structures of protein-ligands complexes shows that both median, minimum, and maximum lengths of the hydrogen bond depend on the interacting partners and are correlated to the hydrogen donor strength. In the fingeRNAt program, we used the single, fixed distance cut-off values for a given interaction type. For example, the default cut-off value for all hydrogen bonds is 3.9 Å, and this distance does not depend on the interacting partners. However, thanks to the plugin-ready architecture of the fingeRNAt, this limitation can be easily circumvented by defining custom hydrogen bond types for atoms/groups of interest and with adjusted distance criteria.

The numbering of residues in the receptor structure must be unique (i.e., there should be no residues with the same number); however, multiple nucleic-acid chains are supported. Also, fingeRNAt currently operates on strictly defined file formats (pdb for nucleic acid receptors and sdf for ligands).

## Datasets

### Non-redundant RNA-small molecule ligand dataset

The non-redundant dataset of complexes of RNA with small molecule ligands was based on the diversified dataset prepared by Philips *et al*. (50). Buffer components and inorganic ions other than metal cations were removed. All complexes were inspected visually, removing or correcting ligands with erroneous chemical structure, missing atoms, or ambiguous stereochemistry. Ionization of the ligands was corrected according to the literature data and was supported by pK_a_ calculations with the Chemicalize platform (95). The final dataset contains 207 RNA-Ligand structures (see S20 Table).

### HIV-1 trans-activation response element activity dataset

To construct a dataset containing information on active and inactive ligands towards human immunodeficiency virus type 1 (HIV-1) trans-activation response element (TAR), an extensive literature search was performed. Information about the chemical structures of molecules and their binding affinities was collected and tabulated together. Ligands, whose binding affinity was not described precisely, or was calculated only on the grounds of cell-based assays, were not included in the database. If the HIV-1 TAR sequence, for which the ligand’s activity is described, differed from the sequence of experimentally solved structure 1UTS (taken arbitrarily) within the binding site or its proximity up to 8Å, the ligand was rejected. Structures of the ligands were normalized, and a set of filters was applied to include only drug-like molecules (molecular weight from 90 to 900 daltons, an octanol-water partition coefficient - SlogP from -7 to 9, number of hydrogen bond acceptors up to 18, number of hydrogen bond donors up to 18, number of rotatable bonds up to 18). Next, the PAINS (Pan-assay interference compounds, (96)) filter was applied to exclude ligands for which observed activity was potentially a result of interference with assays and not the interactions with the RNA. Finally, ligands were assigned into one of two groups - active or inactive. The criterion was the activity (expressed as K_D_, IC_50_, MIC, or K_i_), and ligands with an activity parameter below 300 μM were classified as active. Ligands with the activity parameter higher than 300 μM, or described by the authors as “inactive”, were classified as inactive. Afterward, to remove structurally similar compounds, both active and inactive ligands were clustered using the *k*-medoids algorithm, and for each cluster, a representative ligand was selected. Data processing and analysis were performed using the KNIME 4.0.1 analytics platform (97). As expected, the number of inactive compounds in the dataset was low (10 compounds). To simulate the content of the real chemical library, where the percentage of active molecules is low, additional putative inactive ligands were generated using the DUD-E methodology (98). The final dataset consisted of 30 active and 1478 inactive compounds.

### SIFt-based structure similarity assessment

The goal of the RNA-Puzzles 23 was to predict the structure of the Mango-III (A10U) aptamer bound to TO1-Biotin (PDB code: 6E8U, ligand structure: S10A Fig). RMSD (root-mean-square deviation) values for submitted ligands were calculated for aligned RNA (PyMOL 2.5.0, function align with cycles=0 parameter) using the LigRMSD web server with FlexibleMatch option (allowing the matching of a pair of different atom and bond types when the structure of the submitted ligand was slightly different from the reference ligand; https://ligrmsd.appsbio.utalca.cl/, (75)). As some teams submitted only the fragment of the ligand, for calculation of RMSD, we used the maximum common scaffold present in all submissions - Thiazole Orange *N-*acetamide (SMILES: C[N+]1=CC=C(CC2=[N+](CC(N)=O)C3=CC=CC=C3S2)C2=CC=CC=C12, S10B Fig). RMSD and INF (interaction network fidelity) values for the RNA models were provided by the RNA-Puzzles’ organizers.

Molecular docking was performed with the rDock (version 2013.1; 400 poses were generated); molecular fingerprints were calculated for docked poses with RDKit hydrogens added (-addH rdkit option of the fingeRNAt). RMSD was calculated for the complete ligand structure. The similarity of fingerprints was expressed as the Tanimoto coefficient (expressing the fraction of interactions common for the reference structure and a model) and the Tversky distance (with commonly used values of *α*=1 and *β*=0 and thus representing the fraction of the interactions in the reference structure which are recapitulated in a model) (72).

## Computational protocols

### Calculations of fingerprints

In all case studies presented in this work, the FULL variant of interaction fingerprint was used with the -detail parameter, and its outputs were used for further calculations.

### Molecular docking

The rDock docking program (version 2013.1) was used with dock_solv desolvation potential and a docking radius set to 10 Å (47). For docking of small molecule ligands to HIV-1 TAR RNA, the 1UTS structure was used (99); macromolecules were preprocessed with the Chimera dockprep pipeline (100,101). Fingerprint type FULL was calculated for the best-scored pose of small molecule ligands docked to HIV-1 TAR RNA.

### Analysis of fingerprints

Fingerprint analyses and visualization were performed in the jupyter notebook with the Python 3 kernel. Principal Component Analysis (PCA) calculation and *k*-means clustering were performed using the scikit-learn Python module (102). Optimal clustering parameters were determined by probing a series of cluster numbers and evaluating the quality of clustering with scores: silhouette score, Calinski-Harabasz score, and Davies-Bouldin score (see: S14 Table). Statistical tests were performed in SciPy using the *t*-test for the means of two independent samples of scores, and the two-tailed *p*-value is reported (103).

### Software availability

The fingeRNAt program is freely available and distributed under the open-source GPL-3.0 License. It can be downloaded, along with a manual, collection of helper utilities, and sample data from https://github.com/n-szulc/fingeRNAt/. The program was extensively tested on Python 3.6, 3.7, 3.8, and 3.9 under Ubuntu Linux (18.04, 20.04, and 21.10) and macOS (macOS Catalina 10.15).

## ACKNOWLEDGMENTS

We thank Dr. Marcin Magnus and Dr. Zhichao Miao for providing RNA-Puzzles’ datasets. This research was carried out in part with the support of the Interdisciplinary Centre for Mathematical and Computational Modelling (ICM) University of Warsaw under computational allocation no GB76-20 to F.S.

## SUPPORTING INFORMATION CAPTIONS

**S1 Table. Comparison of the features of the fingeRNAt software (this manuscript) and similar programs (Arpeggio, PLIP2021 and ProLIF)**.

**S2 Table. The total number of interactions detected for RNA-ligand complexes and the number of RNA-ligand complexes with at least one occurrence of a given interaction**.

**S1 Fig. The number of distinct RNA residues forming a given interaction**.

**S2 Fig. Percentage of structures with ligands having at least one of given molecular features**.

**S3 Table. Statistics of complexes and detected Pi-stacking interactions in the RNA-ligand dataset for ligands with or without at least one aromatic ring**.

**S4 Table. Statistics of complexes and detected hydrogen bonds in the RNA-ligand dataset for ligands with or without at least one hydrogen bond donor and acceptor**.

**S5 Table. Statistics of complexes and detected cation-anion interactions in the RNA-ligand dataset for ligands with or without at least one charged atom**.

**S3 Fig. Histograms of bond lengths for non-covalent interactions in a dataset of experimentally solved RNA-ligand structures**.

**S6 Table. Statistics of the lengths for observed non-covalent interactions in a dataset of experimentally solved RNA-ligand structures**.

**S7 Table. Statistics of hydrogen bonds formed by different RNA atoms**.

**S8 Table. Statistics of lipophilic interactions formed by different RNA atoms**.

**S9 Table. Statistics of Pi-anion interactions formed by different RNA groups (when RNA is an anion acceptor) and atoms (where RNA is an anion donor)**.

**S10 Table. Statistics of halogen bonds formed by different RNA atoms**.

**S11 Table. Statistics of all types of interactions formed by different RNA atoms and groups**.

**S12 Table. Linear least-squares regression R**^**2**^ **values calculated for parameters of structures from the RNA-Puzzles collective experiment and redocking experiment**.

**S13 Table. Quality of models submitted to the RNA-Puzzles competition round 23. S4 Fig. The RNA-Puzzles round 23 solution and the best scored model**.

**S5 Fig. Tanimoto coefficient and Tversky similarity of interaction fingerprints calculated for complexes submitted to the RNA-Puzzles round 23**.

**S6 Fig. Relationship between RMSD of ligand and various measures of the interaction fingerprint similarity/distance, calculated for the data from the redocking experiment**.

**S7 Fig. Similarity vs. RMSD of ligands for structures from the RNA-Puzzles collective experiment round 23, calculated for three different resolutions of fingerprints (SIMPLE, PBS, and FULL)**.

**S14 Table. Silhouette score, Calinski-Harabasz score, and Davies-Bouldin score calculated for a various number of clusters, for all-interactions dataset and the one with lipophilic interactions removed**.

**S15 Table. Composition of the input dataset and clusters (number of active and inactive compounds, and percentage of active compounds in a given cluster), and *p*-values for comparing the ratio of active compounds in the input dataset and in a given cluster**.

**S8 Fig. Interaction fingerprints calculated for data from molecular docking of a set of active and inactive compounds to HIV-1 TAR RNA structure 1UTS mapped on the two-dimensional space, with lipophilic interactions removed**.

**S9 Fig. FULL fingerprint calculation times for docking poses of small molecule ligands of various sizes (guanidine, ibuprofen, and sildenafil) to guanidine III riboswitch (RNA with 39 residues)**.

**S16 Table. Mean FULL fingerprint calculation times (in seconds, the average from 10 experiments) of docking poses of small molecule ligands of various sizes (guanidine, ibuprofen, and sildenafil) to guanidine III riboswitch (RNA with 39 residues)**.

**S17 Table. Definitions of the criteria of nine non-covalent interactions calculated by the fingeRNAt.py**.

**S18 Table. Parameters accepted by the fingeRNAt.py**.

**S19 Table. Summary of detected chemical properties and external modules used in the fingeRNAt.py**.

**S20 Table. List of structures used to derive statistics on RNA-small molecule ligand interactions**.

**S10 Fig. Structures of small molecule ligands used in modeling**.

